# Numerical calculation of the light propagation in tapered optical fibers for optical neural interfaces

**DOI:** 10.1101/2021.02.08.430223

**Authors:** Rosa Mach-Batlle, Marco Pisanello, Filippo Pisano, Massimo De Vittorio, Ferruccio Pisanello, Cristian Ciracì

## Abstract

As implantable optical systems recently enabled new approaches to study the brain with optical radiations, tapered optical fibers emerged as promising implantable waveguides to deliver and collect light from sub-cortical structures of the mouse brain. They rely on a specific feature of multimodal fiber optics: as the waveguide narrows, the number of guided modes decreases and the radiation can gradually couple with the environment. This happens along a taper segment whose length can be tailored to match with the depth of functional structures of the mouse brain, and can extend for a few millimeters. This anatomical requirement results in optical systems which have an active area that is very long compared to the wavelength of the light they guide and their behavior is typically estimated by ray tracing simulations, because finite element methods are too computationally demanding. Here we present a computational technique that exploits the beam-envelope method and the cylindrical symmetry of the fibers to provide an efficient and exact calculation of the electric field along the fibers, which may enable the design of neural interfaces optimized to meet different goals.

## I. Introduction

Optogenetics is a powerful technique that uses light to control the electrical activity of neurons genetically modified to be light sensitive [1, 2]. In the last decade, this tool has been able to revolutionize the field of neuroscience because it allows for cell-type specificity, it provides very high spatial and temporal precision, and it can target multiple areas of the brain with multiple wavelengths [3]. Moreover, some of the tools used in optogenetics to deliver light to the brain, including optical fibers, have been shown to allow the recording of the optical activity of different indicators and molecules [4, 5]. The design of optical neural interfaces able to deliver light in a controlled manner in deep brain regions is an essential ingredient to achieve an optimal interaction with neurons and exploit the full potential of optogenetics [6]. While flat cleaved fiber optics enable an efficient and controllable targeting of shallow brain regions, the targeting of elongated structures (beyond 1 mm) is prohibited by tissue scattering and absorption [7, 8, 9]. A promising candidate for developing a new generation of neural interfaces able to target deep regions extending up to 3 mm are tapered optical fibers, which can perform both homogeneous light delivery and dynamically-controlled spatially-restricted illumination of the brain [10, 11]. To control how light reaches the brain one can tune the input light distribution and choose the appropriate fiber numerical aperture and tapering angle. Indeed, the number of supported modes decreases as the waveguide narrows, and non-guided modes can couple with the environment. It has been demonstrated that the taper operates a mode-division demultiplexing of guided light and this peculiar property has been exploited to obtain depth-resolved optical neural interfaces. In practice, the smaller the input angle of incident beam with respect to the fiber axis, the deeper the region where the tapered fiber delivers light, a feature that offers the possibility of targeting different brain areas along the implant axis with reduced invasiveness [12, 13, 14, 15, 16].

In order to match the spatial extent of elongated brain regions and achieve an effective mode-division demultiplexing, the length of the tapered region must be of the order of some millimeters, much larger than the wavelength range typically employed in optogenetics (450-600 nm). As this makes traditional finite element calculations computationally very demanding, the power that is guided and delivered to the brain is usually estimated using ray tracing simulations [10, 11, 12, 13]. These calculations help predict at what distance from the fiber tip the light is delivered but they do not provide an estimation of the electric and the magnetic field distribution in the targeted region, since they fail to describe interference and diffraction phenomena. This fact limits our ability to optimize the delivery and collection of light, and makes it difficult to design new optical neural interfaces with different functionalities.

One of the most recent approaches to expand the functionalities of optical fibers consists in modifying the end-face or the tip of tapered optical fibers to include different types of structures that interact with the incident light in intriguing ways [17, 18]. For example, several studies demonstrate that the use of plasmonic structures surrounding the fiber could help confine and concentrate light at the nanoscale [19, 20, 21, 22]. Also, structures made of high-index dielectrics [23] or metals [24] have been shown to enable sub-diffraction light confinement. These studies usually employ single-mode optical fibers and therefore do not consider the mode-division demultiplexing of guided light in tapered optical fibers. If one had access to the field distributions along the taper as well as to the evanescent field around it, the design of all these structures could be optimized in order to further enhance the light-brain interaction and eventually lead to optimal depth-resolved optical neural interfaces.

In this work we propose a strategy for calculating the electric and the magnetic fields distribution in long tapered fibers (TFs) using a computational method which is much more cost efficient than traditional full-wave finite element calculations. First, we exploit the cylindrical symmetry of the modes that the fiber can guide in such a way that the fields are not calculated from a three-dimensional (3D) model but from the superposition of a finite set of two-dimensional (2D) axis-symmetric models. Second, we assume that the variation of the amplitude of the fields along the fiber axis, which is due to the light reflections at the taper interfaces, are much slower than the oscillations due to the phase propagation. In this way, we do not need to calculate the field oscillations along the taper axis, which would require a fine mesh able to resolve the wavelength of the incident light. Instead, we compute the variations of a slowly varying envelope function that can be evaluated using a much coarser mesh and, therefore, significantly reduced computation resources. To demonstrate the validity of the proposed methodology, we compare the obtained numerical results with experimental results. In particular, we compare the light delivery depth at different input angles for two different fibers.

## II. Calculation of the incident field distribution

In this work, we consider multi-mode fibers with both core and cladding tapered as depicted in Fig. 1a [13]. As the light propagates through the taper, guided modes supported at a given section morph or couple into new radiative modes. Because of this change in the nature and the number of modes that need to be considered, such problem cannot be solved in the so-called adiabatic regime [25]. As a first step for calculating the field propagation in long TFs, one needs to obtain the field distribution at the input fiber facet, which will depend on the fiber geometry and materials, as well as on the excitation. The input field then will be propagated along the taper through the method described in the next section.

**Fig. 1.**
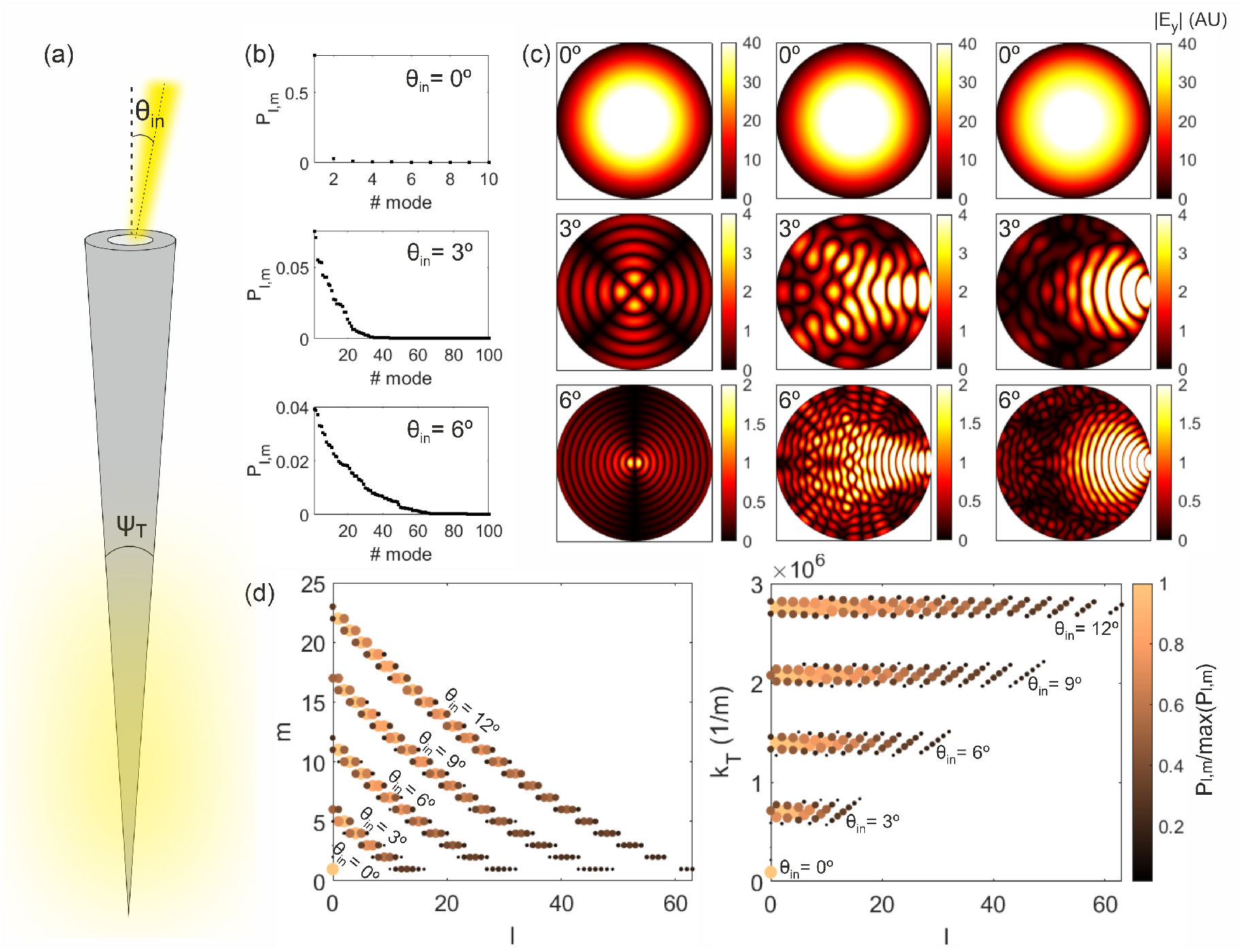
(a) Sketch of a step-index tapered optical fiber. (b) Fraction of the power guided by the modes supported by a *NA* = 0.22 fiber of core diameter *a* = 50 *µ*m when excited by a *y*−polarized Gaussian beam of wavelength *λ*_0_ = 473 nm, waist radius 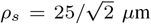 and incident angle *θ*_in_ = 0°, *θ*_in_ = 3°, and *θ*_in_ = 6°. The modes are ordered according to the amount of power they guide, so for each *θ*_in_ the mode number 1 is the one guiding the largest power. (c) Absolute value of the component *E*_*y*_ of the electric field at the input core facet of the TF in (b) for *θ*_in_ = 0° (top), *θ*_in_ = 3° (middle), and *θ*_in_ = 6° (bottom) considering the mode guiding the largest power (left) and the superposition of the modes guiding at least a 50% (middle) and a 95% (right) of the total guided power. (d) Numbers *l* and *m* and transverse component of the propagation constant *k*_T_ of the modes guiding the 95% of the total power guided by the fiber for *θ*_in_: 0°, 3°, 6°, 9°, and 12°. The color and size of the markers indicate the power of each mode normalized to the power of the mode guiding the largest power for each *θ*_in_.

Consider a step-index optical fiber consisting of a cylindrical dielectric core of refractive index *n*_1_ surrounded by a cylindrical cladding of refractive index *n*_2_ slightly lower than *n*_1_ in the presence of a light beam impinging with an angle *θ*_in_, as depicted in Fig. 1a. When the incident light reaches the fiber entrance, it can excite one or multiple modes in the fiber. The number of modes that a fiber is able to guide is determined by the normalized frequency of the fiber, *V*, which is defined as *V* = *πa/λ*_0_*NA*, where 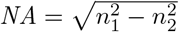 is the fiber numerical aperture, *a* is the fiber core diameter, and *λ*_0_ is the wavelength of the incident light. Thus, given a certain wavelength of the incident light, fibers with larger cores or with larger numerical apertures are able to guide a larger number of modes.

In the weakly guiding approximation (*n*_2_ ≈*n*_1_), the longitudinal components of the electric and the magnetic field (*E*_*z*_ and *H*_*z*_) are negligible. In this case, the fiber guides a set of linearly polarized modes, which are identified by the indexes *l* and *m* and denoted as *LP* modes [26, 27]. For each (*l, m*) there are four degenerate modes sharing the same propagation constant *β*_*l,m*_. For a given light source impinging on the input fiber facet one can obtain the power carried by each of the excited modes as the overlap integral between the input radiation and the modes at the fiber input. In particular, the amplitude of the excited (*l, m*) mode can be calculated as [10]:

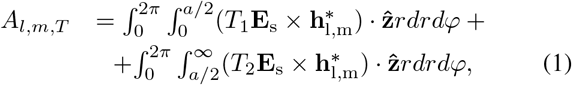

where *T*_1_ and *T*_2_ are the transmission Fresnel coefficients, **E**_s_ is the transverse component of the electric field incident on the fiber input facet, and **e**_l,m_ and 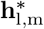 are the transverse electric and magnetic field profiles, respectively, of a given degenerated (*l, m*) mode (normalized such that 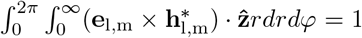. The ratio of the energy carried by each mode then is *P*_*l,m*_ = | *A*_*l,m,T*_ | ^2^. The field profiles of the LP modes and their propagation constants are well known and they can be found in [27], for example. The superposition of all the modes will result in the field distribution at the fiber input facet.

In our calculations, we consider a linearly polarized Gaussian beam forming an angle *θ*_in_ with the fiber axis. The numerical aperture of the fiber limits the input angles that can be used to couple light into the fiber to *θ*_a_ = sin^−1^ NA [26]. For a fiber with NA = 0.22, for example, *θ*_a_ is found to be 12.7°; for this reason experimental data is only reported for *θ*_in_¡12.7 [11]. In this range of *θ*_in_, the transverse component of the electric field incident in the fiber input facet can be written as:

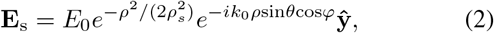

where *k*_0_ is the wave number in free space, *ρ*_*s*_ is the Gaussian beam waist radius and *E*_0_ is a normalization factor 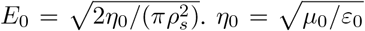 is the impedance of free space, with *µ*_0_ and *ε*_0_ being the vacuum permeability and permittivity, respectively. Throughout this work, we consider the incident light wavelength being *λ*_0_ = 473 nm, one of the most commonly used in optogenetics (it corresponds to the activation wavelength of the protein Channelrhodopsin-2, ChR2); nonetheless, all the parameters we use in the simulations can be tuned to meet the requirements of different optogenetics tools [28].

Figure 1b shows the fraction of the power guided by the modes supported by a fiber with NA = 0.22 and core diameter *a* = 50 *µ*m for three different incident angles of a linearly polarized Gaussian beam of waist radius 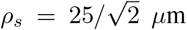 calculated using Eqs. (1) and (2). The modes are sorted in such a way that mode 1 is the one guiding the largest power and that the power decreases as the mode number increases. It can be seen that the larger the incident angle, the larger the number of modes that carry an appreciable proportion of the incident light power.

The electric field distribution at the input fiber facet can be obtained as the sum of the electric field created by all the modes supported by the fiber, each of them weighted according to the power they guide for each incident angle. Figure 1c illustrates the field distribution at the core input facet of the fiber calculated taking into account only the mode guiding the highest power (left) and the superposition of all the modes guiding a 50% (middle) and a 95% (right) of the total guided power for the three incident angles in Fig. 1b. These examples demonstrate that for incident angles different from 0° the field distribution will strongly depend on the number of modes taken into account in the calculations.

Finally, Fig. 1d illustrates that the larger the input angle, the wider the range of mode numbers *l* and *m* that are excited with a significant portion of the power. Larger input angles *θ*_in_ excite modes with larger values of *l* and *m*: for *θ*_in_ = 3°, for example, *l* ranges from 0 to 16 and *m* from 1 to 6, while for *θ*_in_ = 12° *l* ranges from 0 to 63 and *m* from 1 to 23. The right panel shows that the value of the transverse component of the propagation constant 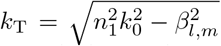 associated to the guided modes increases with the incident angle. Also, it can be seen that for each incident angle the power of modes with higher values of *l* is lower than that of modes with lower *l* values.

## III. Beam envelope method for calculating the field propagation in long optical fibers

Once the field at the fiber input is determined, one can proceed to the calculation of the electric and magnetic field distribution along the taper in order to obtain the fields that will reach the outer surface of the TF and, therefore, will be delivered to a targeted region of the brain. To this end, one could think of creating a 3D model of the long TF with the calculated input field distribution at the fiber entrance. However, this model would require an extremely thin mesh compared to the dimensions of the fiber, resulting in very high computational demands. Indeed, standard taper lengths are >1 mm, which is more than three orders of magnitude longer than the wavelength *λ*_0_ = 473 nm. Since one usually needs at least 5 discretization points per wavelength (in the medium) to resolve a wave oscillation, the calculations would require a very fine mesh in the axial fiber direction (hereafter referred to as *z*−, see axes definition in Fig. 2a). Moreover, one would also need to discretize a large 3D space with a mesh sufficiently thin in the *x*- and *y*-directions to resolve all the light reflections along the taper interfaces. Thus, some strategies for reducing the computation requirements are necessary.

**Fig. 2.**
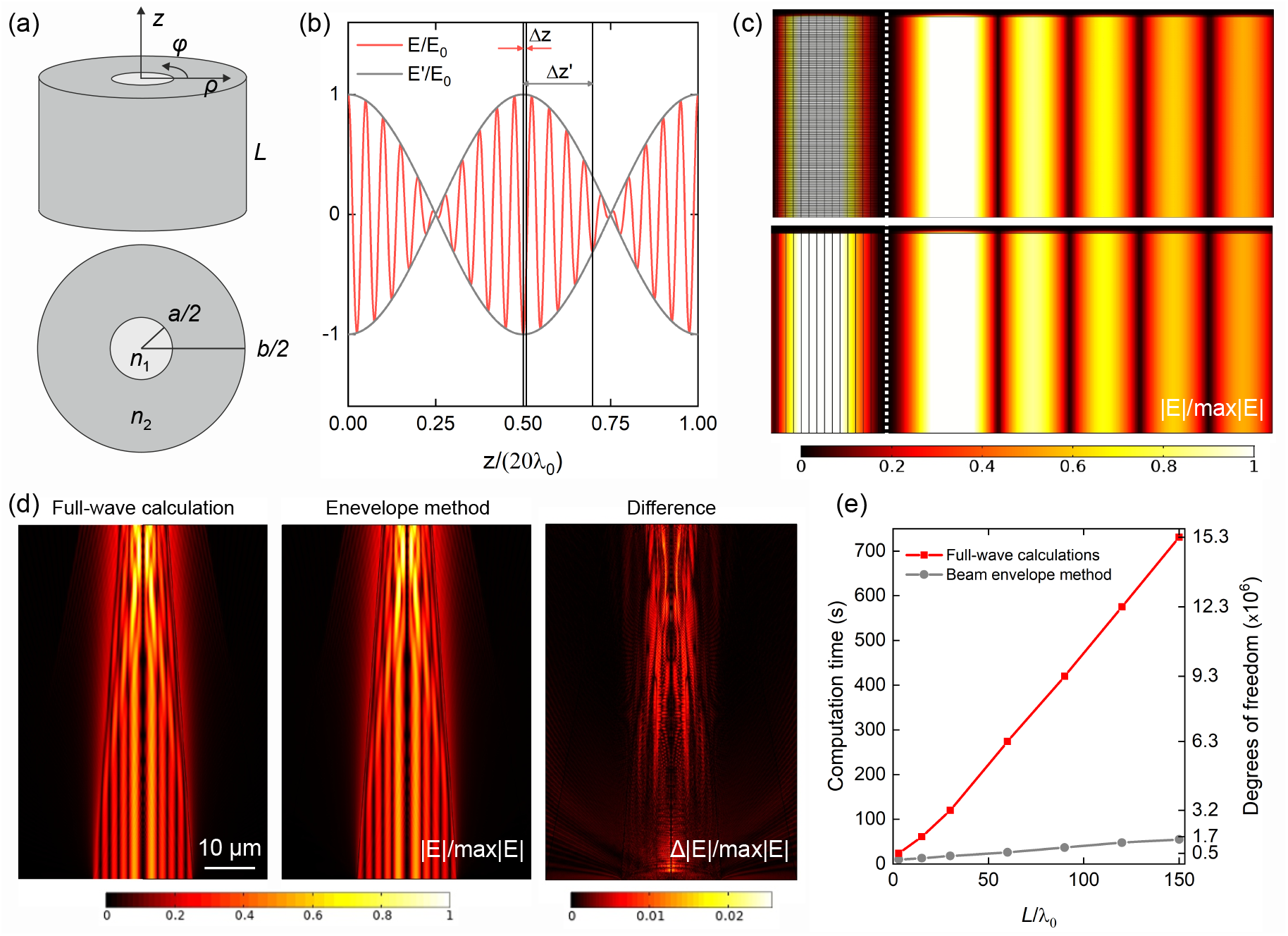
(a) Sketch of a segment of length *L* of a step-index optical fiber. (b) Illustration of the propagation of the electric field **E** and its envelope **E’** normalized to the incident field amplitude *E*_0_ showing the required minimum mesh elements size to resolve both waves (5 nodes per wavelength). (c) Electric field in the core of a segment of length *L* = 30*λ*_0_*/n*_1_ of a step-index optical fiber of *NA* = 0.22 and core diameter *a* = 50 *µ*m when excited with the mode *l* = 2, and *m* = 5. The first panel corresponds to the calculation with a traditional finite element method and the second panel to the calculation with the envelope method. For comparison, the mesh used in these calculations is shown in the left side of the plots. The dotted white line indicates the fiber axis. (d) Evolution of the mode *l* = 2 and *m* = 5 along a fraction of length *L* = 150*λ*_0_ of a TF of *NA* = 0.22 and tapering angle *ψ*_T_ = 20° obtained from a full-wave calculation (left panel), using the beam envelope method (central panel), and the difference between them (right panel). (e) Comparison between the computation time and the degrees of freedom required to calculate the electric field propagation in different fractions of the TF shown in (d) for the traditional full-wave finite element and the envelope method calculations. Simulations were performed on a workstation equipped with two 12-cores CPUs (Intel Xeon 4116 2.1 2400MHz) and 192 (6×32) GB of RAM.

To this aim, the 3D problem can be converted into a set of 2D problems. Even though the total field distribution at the fiber entrance is in general not axisymmetric due for example to the asymmetry of the excitation field, as well as the polarization (Fig. 1c), full-vector fields can be expanded as a sum of cylindrical harmonics with respect to the azimuthal variable *φ* as:

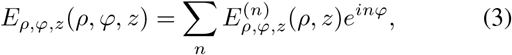

with *n*∈ℤ being the azimuthal mode number. Because the geometry is *φ*-independent (i.e., axisymmetric), each cylindrical harmonic propagates independently satisfying the following equation:

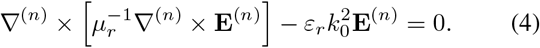

where the operator ∇^(*n*)^ × is obtained from the corresponding standard operator in cylindrical coordinates with the following substitution 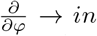. The full 3D problem then is reduced to a set of 2D problems corresponding to different values of *n* (see Refs. 29 and 30 for more details).

Due to the nature of our problem, we could further reduce the computational cost by leveraging the knowledge of the input photon momentum. Consider an electromagnetic wave with electric field **E**(*ρ, φ, z*) propagating along a fiber with propagation constant *β*. If there is no change in the wave momentum, i.e. *β* is unchanged, the electric field can be simply written as **E**′(*ρ, φ*)*e*^−*iβz*^, where **E**′is constant with respect to the propagation direction *z*. However, when the wave enters the tapering region, the slightly tilted boundaries are going to modify the photon momentum such that the new propagation constant becomes *β*′= *β* + *δβ*. Since the tapering angle is small (*ψ*_*T*_ ≤ 3.7°) and since the input mode will couple to modes with very similar propagation constants as the fiber shrinks, we expect *δβ*≪*β* (although this is not a strict requirement as we will show later), and the electric field becomes **E**′(*ρ, φ*)*e*^−*iδβz*^*e*^−*iβz*^. Because *δβ* is unknown and is changing along the propagation, we can incorporate it into the expression of the electric field as **E**′(*ρ, φ, z*)*e*^−*iβz*^, where the amplitude of **E**′is slowly varying along *z*, as illustrated in Fig. 2b. Since **E**′may vary much more slowly in the *z*-direction than **E**, it can be resolved by a much coarser mesh and, as a result, it will require much lower computational resources (mesh size illustrated in Fig. 2b).

In order to solve for **E**′ then, one needs to substitute **E** = **E**′*e*^−*iβz*^ in the wave equation. As a results, the curl of the electric field becomes 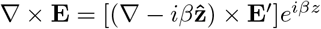. Considering for the field **E**′an expansion similar to Eq. (3) we obtain the following equation:

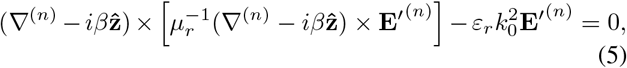

which is solved for different values of *n* in the *ρ*-*z* plane. We have implemented Eq. (5) in the *radio frequency electromagnetic waves* module of Comsol Multiphyics, as described in the Appendix. Note that this equation only represents a recast of the usual wave equations and does not introduce any simplification. The advantage of using such formulation is that we are leveraging the knowledge on the mode momentum to reduce the numerical discretization required to resolve the field oscillations. In fact, if the condition *δβ* ≪ *β* is not satisfied, this would translate in a mesh requirement comparable to the standard full-wave implementation. On the other hand, when the condition is verified this method introduces a relaxation on the mesh requirement along *z* of the order of *β/δβ*.

Finally, we need to determine the excitation field at the input facet and the propagation constant *β*. This can be done by using the analytical solutions of a flat fiber with geometrical parameter corresponding to the input facet of the taper. By means of Eq. (1) we can project the excitation field onto the fiber modes. Naturally, each mode can be easily written in cylindrical coordinates. Assuming that the incident field is polarized in the *y*-direction, the angular and the radial components of the electric field distribution of the mode (*l, m*) with propagation constant *β*_*l,m*_ can be written as:

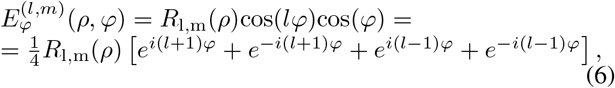

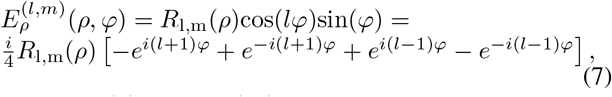

where *R*_l,m_(*ρ*) and cos(*lφ*) are the radial and the angular dependence of the *y* component of the (*l, m*)-mode electric field [26]. For each (*l, m*)-mode we then can calculate its evolution along the taper by solving 4 axisymmetric calculations, with azimuthal mode numbers *n* = ± (*l*±1) (the number of calculations for each mode does actually reduce to 2 since ± *n* azimuthal mode numbers give symmetric results) and taking *β* = *β*_*l,m*_.

It is worth noting that similar envelope methods have been applied in laser plasma accelerators, in which laser pulses are guided along very long channels compared to the laser wavelength [31, 32, 33]. Our approach however differs from the beam propagation method [34, 35] and the parabolic equation method [36], both used to simplify the calculations in long waveguides, because it retains the second order derivatives along *z* and hence does not give an approximation but the exact solution of the field distribution, provided that the envelope oscillations are well resolved. Another similar method well-suited to analyze taper structures is the eigenmode expansion (EME) method [37, 38], which relies on the decomposition of the electromagnetic fields into a basis set of local eigenmodes that exists in the cross-section of the waveguide. The EME however is not very efficient for modeling structures requiring a very large number of modes, and especially for structures that vary continuously along the propagation direction since it requires all the modes to be re-computed at discrete positions [39]. Indeed, a full taper would require an increasing number of radiative modes to be included in the calculation. Our approach, on the other hand, naturally takes into account any relevant mode, since the input mode can freely evolve along the taper.

To verify our envelope method, we compare the total field distribution to the result obtained with full-wave finite element calculations. First, we consider a short segment of length *L* of a flat step-index weakly guiding optical fiber of core and cladding refractive indexes, *n*_1_ = 1.464 and *n*_2_ = 1.447, and core and cladding diameters *a* = 50 *µ*m and *b* = 125 *µ*m, respectively. As a light source, we consider a linearly polarized Gaussian beam forming an angle *θ*_in_ = 3° with the fiber axis. We calculate the evolution of the mode *l* = 2, *m* = 5, which guides the largest portion of the total power guided for this incident angle. The field distribution of this mode at the fiber entrance was shown in Fig. 1c. As shown in Fig. 2c both methods give the same total field distribution along the whole length of the fiber. The meshes are compared on the left side of Fig. 2(c). In both cases we have considered the same number of points in the radial direction and the minimum number of mesh elements required for calculating the field propagation. In the case of the traditional finite element calculations, each wavelength is resolved by 5 elements (in the *z*-direction). In contrast, the envelope method makes it possible to calculate the envelope propagation using a single element for all the length (Fig. 2c, bottom) and, therefore, the mesh elements do not vary with *L*. Thus, while the computation time required to perform a traditional finite element calculation linearly increases with the fiber length, the computation time of the beam-envelope calculations does not depend on the fiber length, it remains constant at 3 s.

This case corresponds to an ideal scenario in which there are no light reflections and the amplitude of the envelope is constant, i.e. it does not oscillate with *z*, and one is able to carry out the envelope-function calculations using a single mesh element. In general, including the cases we will study below, there are reflections in the fiber due to the tapering, for example, which results in oscillations in the electric field envelope and a mesh that resolves the envelope oscillations is needed. In order to verify that our method gives accurate results also when there are reflections in the fiber, we assume that the fiber considered in Fig. 2c is tapered, with a tapering angle *ψ*_T_ = 20°. The field is calculated on a short section of the fiber of length *L* = 150 *λ*_0_ where the input facet core diameter was reduced to *a* = 20 *µ*m so the full-wave finite element calculations can run in a reasonable amount of time. Results in Fig. 2d illustrate that the field distribution obtained with the envelope method looks identical to the field distribution obtained with traditional full-wave calculations: the difference between both calculations is smaller than 2.5%. Finally, Fig. 2e shows the computational time required to calculate the field distribution along different portions of the fiber in (d) with both methods. The computational time increases with the fiber length, but it is much lower when using the envelope function than when using the traditional finite element calculations, specially when considering long fibers. Moreover, the difference between the results calculated using the beam-envelope method and traditional full-wave calculations does not increase with the fiber length. The average error, calculated as the average of the difference between field modulus divided by its peak value (plotted in panel 3 of Fig. 2(d) for the case L = 150*λ*_0_) ranges between 0.05% and 0.3% for all the considered fiber lengths.

## IV. Results and discussion

We can now apply the proposed strategy to calculate the total field distribution in long tapered optical fibers. Since we are interested in comparing our results with experimental data, we focus on the analysis of two different fibers whose behaviour as neural interfaces has already been experimentally characterized [11, 13]. In the following, we assume that the modes excited by the Gaussian beam propagates unaltered through the optical fiber until they reach the tapered region, where their evolution is obtained by solving Eq. (5).

The first fiber is defined by the following parameters: numerical aperture *NA* = 0.22, core and cladding diameters *a* = 50 *µ*m and *b* = 125 *µ*m, respectively, and core and cladding refractive indexes *n*_1_ = 1.464 and *n*_2_ = 1.447, respectively. The second fiber is defined by *NA* = 0.66, *a* = 200 *µ*m, *b* = 230 *µ*m, *n*_1_ = 1.63, and *n*_2_ = 1.49. The fibers are assumed to be surrounded by water, with refractive index *n*_*w*_ = 1.33. The tapering angle (defined in Fig. 1a) is *ψ*_T_ = 2.2° for the *NA* = 0.22 fiber and *ψ*_T_ = 3.7° for the *NA* = 0.66 fiber. The fiber tip is assumed to be cut at a diameter *d*_T_ = 500 nm, which leads to a fiber length, *L*_T_, of 3.242 mm and 3.553 mm for the *NA* = 0.22 and the *NA* = 0.66 fibers, respectively. In all the calculations we will consider a linearly *y*−polarized Gaussian beam of wavelength *λ*_0_ = 473 nm forming an angle *θ*_in_ with the fiber axis. For fibers with *NA* = 0.22 the maximum *θ*_in_ that will be considered is *θ*_in_ = 12°, while for *NA* = 0.66 fibers the incident angle can go up to *θ*_in_ = 33° due to their larger numerical aperture.

To start with, consider an incident angle *θ*_in_ = 0°. It can be obtained from Eqs. (1) and (2) that for this incident angle the 95% of the total power guided by the fiber is carried by only three modes, both for *NA* = 0.22 and *NA* = 0.66 fibers. The modulus of the total electric field distribution in the tip of these fibers is shown in Fig. 3 for these three modes. We only plot here the field at the very tip of the fiber for illustration purposes (the fiber entrance is set at *z* = 0 mm), as in this way it is possible to observe how the field injected by each mode is delivered to the taper surrounding media.

**Fig. 3.**
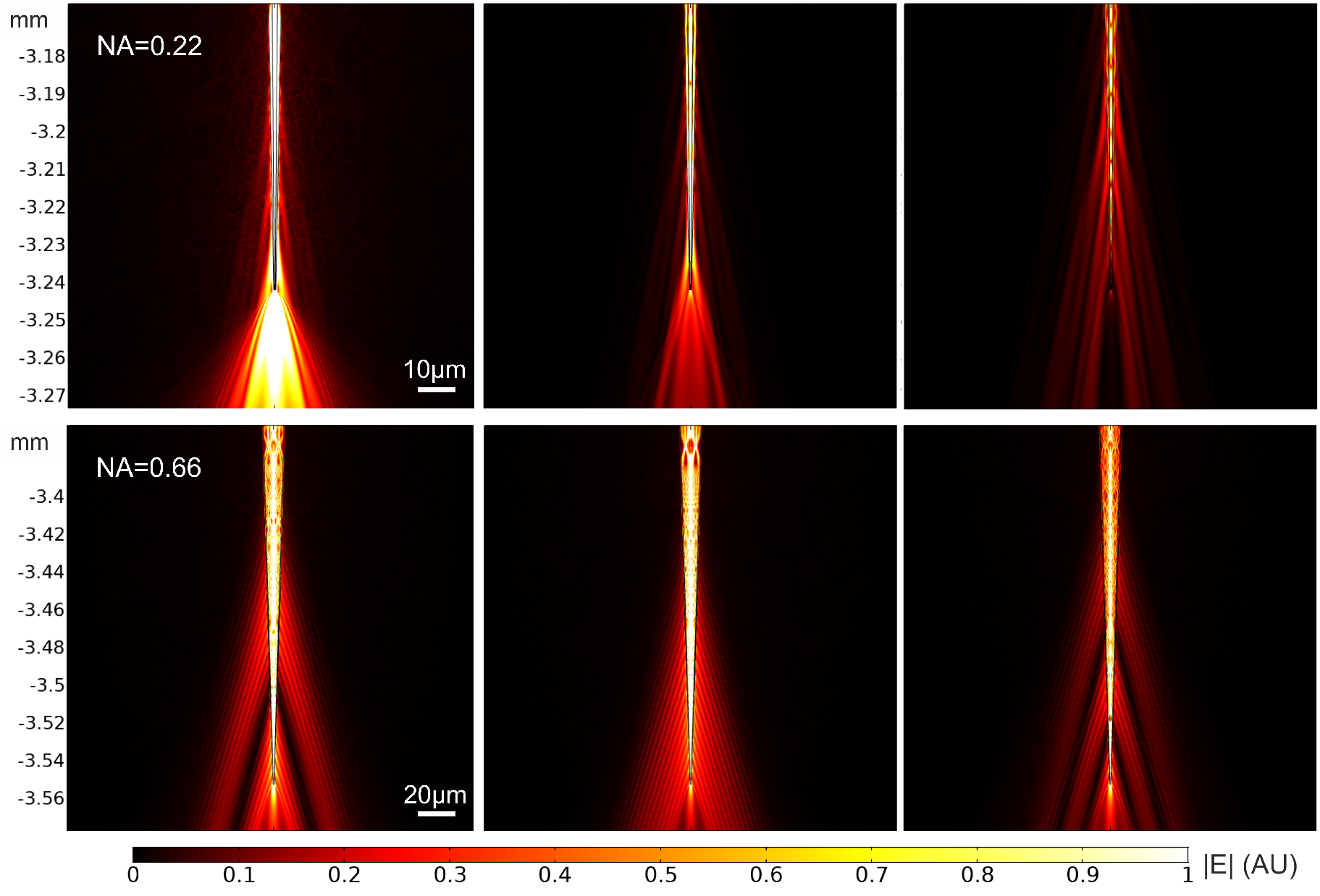
Evolution of the three modes guiding the largest power when the NA = 0.22 (top) and the NA = 0.66 (bottom) fibers are excited with an incident angle *θ*_in_ = 0°.

The number of modes that guide a significant fraction of the power increases with the incident angle, as seen above (Fig. 1b), which makes it difficult to illustrate the evolution of all the excited modes for the other *θ*_in_. However, one can still compare the total field associated to the mode guiding the largest fraction of the power for different *θ*_in_. Such fields are shown in Fig. 4, using intervals of 2° for the *NA* = 0.22 fiber and 5° for the *NA* = 0.66 fiber. Figure 4 shows that for both numerical apertures low input angles generate the light delivery close to the tip, while increasing *θ*_in_ moves the emission farther from it. This result is in agreement with the theoretical and the experimental results reported in the literature [10, 11, 13].

**Fig. 4.**
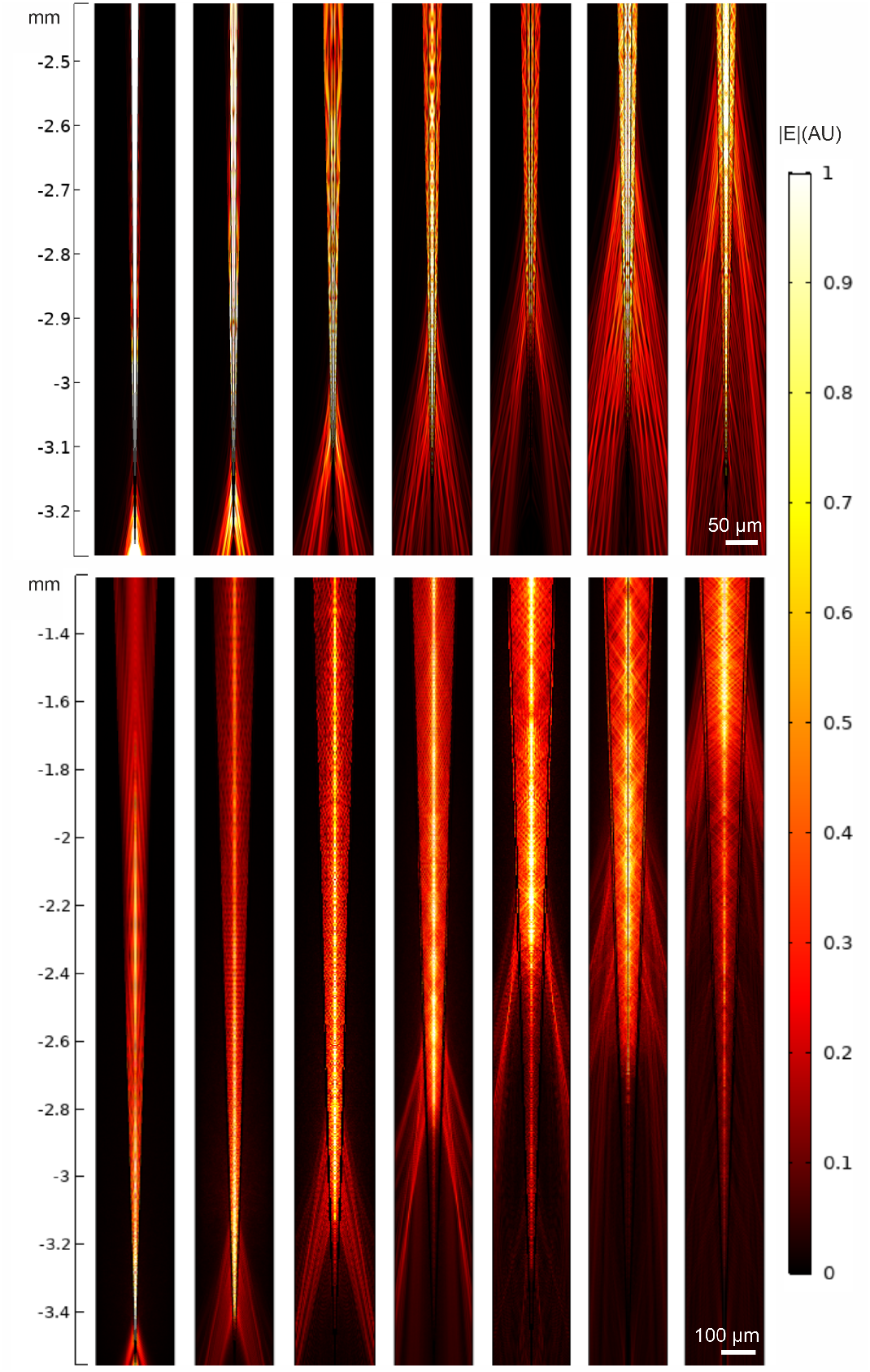
Evolution through the tapered region of the modes guiding the largest power when the NA = 0.22 (top panels) and the NA = 0.66 (bottom panels) fibers are excited with incident angles (from left to right) *θ*_in_ = 0°, 2°, 4°, 6°, 8°, 10°, and 12°, and *θ*_in_ = 0°, 5°, 10°, 15°, 20°, 25°, and 30°, respectively.

The light delivery to the surrounding media can be further characterized by analyzing how the delivery distance from the tip depends on the light incident angle. Figure 5a shows the profile of the square of the electric field, which is proportional to the light intensity, along a line parallel to the external surface of the TF for all the input modes as in Fig. 4. The total field is measured in the surrounding media, at 0.5 *µ*m from the fiber surface (moving in the radial direction). From these intensity profiles one can calculate the starting emission diameter, which is found as the fiber diameter at which the modulus of the intensity is a 90% of half its peak value, and the centroid distance from the tip, defined as the distance at which the integral of the intensity from the tip to that point corresponds to more than 50% of the same integral along the whole taper surface, from the tip to the input facet of the fiber. Both the starting emission diameter and the centroid distance from the tip are found to increase linearly with the incident angle (Fig. 5b). This is because the transverse component of the propagation constant of the guided modes, *k*_T_, which defines the fiber diameter where the modes become evanescent and, thus, are not guided anymore, increases linearly with *θ*_in_ (Fig. 5c).

**Fig. 5.**
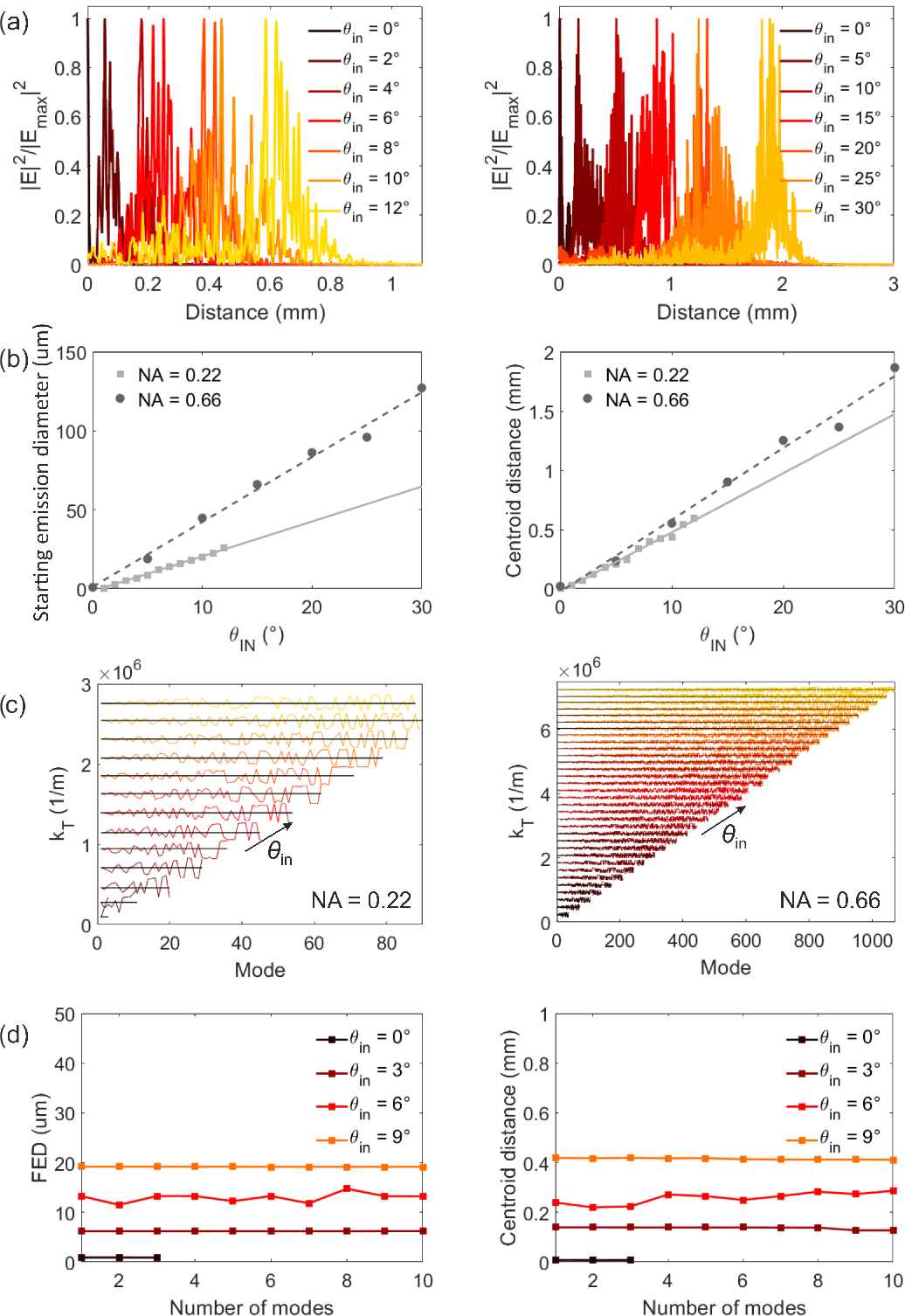
(a) Square of the modulus of the total electric field |*E* |^2^ at 0.5 *µ*m from the fiber surface as a function of the distance from the tip for different incident angles for the *NA* = 0.22 fiber (left) and the *NA* = 0.66 fiber (right). For each incident angle, results correspond to the mode guiding the largest fraction of the total guided power and are normalized to the maximum square of the modulus of the electric field, |*E*_max_| ^2^, for that angle. (b) Starting emission diameter and centroid distance from the tip as a function of the incident angle. The points correspond to the results obtained from the lines in (a) and the lines give the linear fitting of the data. (c) Transverse component of the propagation constant of the modes guiding a 95% of the total guided power sorted in such a way that mode 1 corresponds to the mode guiding the largest fraction of the power for each *θ*_in_. Each color corresponds to a different input angle. (d) Starting emission diameter and centroid distance from the tip for different incident angles as a function of the number of modes that are computed.

In order to compare these results to the experimental data reported in previous works[11, 13], we first need to verify that the starting emission diameter and the centroid distance calculated assuming just one mode for each *θ*_in_ are a good approximation to the results we would obtain if all the modes that are excited for each *θ*_in_ were taken into account. To this end, we calculate the starting emission diameter and the centroid distance from the tip assuming the superposition of different number of modes for different incident angles, from a single mode to ten modes. Since the total field distribution corresponding to the superposition of each mode’s evaluations is not axisymmetric (see Fig. 1c), the field profiles have been obtained by averaging the field distribution along *φ* (at 0.5 *µ*m from the fiber surface, as above). As shown in Fig. 5d the results for the different number of input modes are approximately the same for all the considered input angles: a single input mode seems to be enough to estimate the starting emission diameter and the centroid distance from the tip. This can be understood from the plots in Fig. 5c. Since all the modes that are excited for a given *θ*_in_ have a very similar propagation constant, which oscillates around the propagation constant of the mode guided with the largest power for each *θ*_in_, they all become evanescent at a similar distance from the fiber tip. Therefore, the starting emission diameter and the distance from the tip must be approximately the same for all the modes excited with the same *θ*_in_. This is illustrated in the example in Fig. 3, as the starting emission diameter of the three modes that are guided for each fiber (all three excited with *θ*_in_ = 0°) is approximately the same.

Finally, we can compare the centroid distance from the tip obtained with the numerical calculations to the values obtained from the data reported in Refs. [11] and [13] for the TFs with NA = 0.22 and NA = 0.66, respectively. Experimental light emission profiles were measured from the fluorescence intensity generated by a TF immersed in an homogeneous bath (PBS:Fluorescein 30 *µ*M). Using a fast laser scanning system, the laser input angle *θ*_in_ was controlled to launch modes with a defined transverse component *k*_T_ in a patch cord butt-coupled to the TF in exam. Briefly, a CW laser beam (473 nm, Ciel, Laser Quantum) was focused on a galvanometric mirror that deflects the incoming light at an angle proportional to its driving voltage. The laser beam deflected by the mirror is collected and collimated by a first aspheric lens (focal length 100 mm) and focused on the patch cord input core with a second aspheric lens (focal length 32 mm). The fluorescent light generated by the fiber emission is imaged on a sCMOS camera, synchronized with the scanning mirror, that produces images with a discretization of 0.522 px*/µ*m. This system allows us to image the light emission profiles generated by input angles covering the full angular acceptance of the two fibers, with steps of 0.5°.

For a better comparison with the experimental data, instead of considering a single field profile as in Fig. 5a, we average the total electric field along a set of 21 parallel lines. The first line is set at the fiber surface and the last one at a distance of 2 *µ*m from the surface, so the spacing between lines is 0.1 *µ*m. For each incident angle, we obtain the centroid of the emission region as 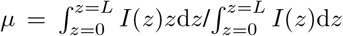, where *L* is the length of the fiber [13], and the standard deviation of the data as 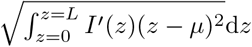, where *I*′(*z*) is 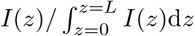.Results are shown in Fig. 6. For the experimental data we have calculated the centroid distance from the tip taking into account the data above the peak half prominence [13]; this avoids the influence of fluorescence tails excited in the medium by the light emitted from the active region. Notice that, since we do not consider data beyond the tip, the centroid we obtain does not always correspond to the point where the field is maximum. This is evident in Fig. 6a for the *NA* = 0.22 fiber with an input angle of 0Â °: the centroid distance from the tip obtained from the experimental data (the profiles are shown in the inset) is found to be 0.13 mm, but looking at the profile in the inset one sees that the maximum of the intensity is at 0 mm.

**Fig. 6.**
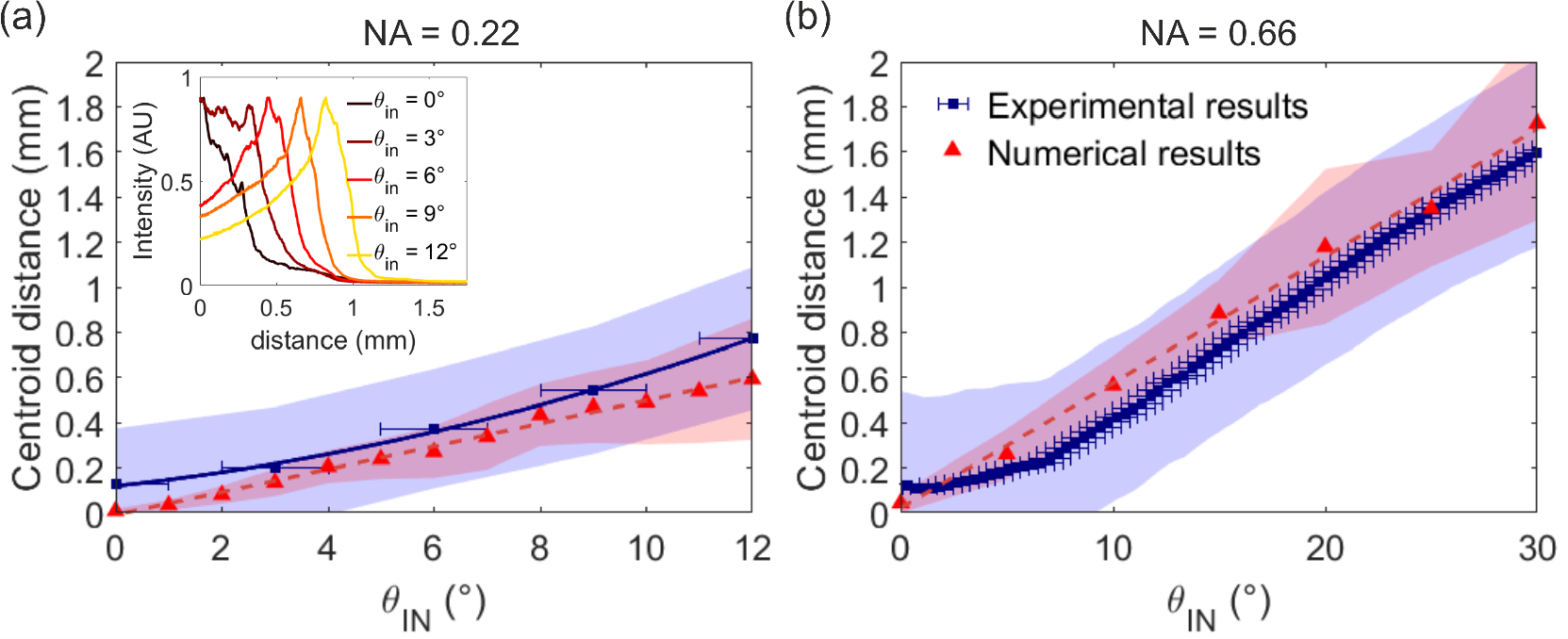
Centroid distance from the tip for the (a) *NA* = 0.22 and (b) *NA* = 0.66 tapered fibers calculated from the intensity profiles obtained from the experimental data (blue squares) and from the numerical calculations (red triangles). The solid and dashed lines are guides for the eye. The experimental results have a convergent input beam of ±1°, used to report the horizontal error bars. The blue and the red shadowed regions are bounded by the centroid distance from the tip ± the standard deviation, which gives an idea of the emission region. The inset in (a) shows the experimental intensity profile along the surface of the NA=0.22 fiber for each incident angle.

Figure 6 shows that the numerical results exhibit a similar trend to that of the experimental results: the centroid distance from the tip increases approximately linearly with the incident angle with a relatively good agreement. For low input angles, and in particular for *θ*_in_ = 0°, the theoretical centroid is close to the fiber tip and has a lower standard deviation, which differs from the experimental results, which exhibit an approximately constant standard deviation for all the input angles. For example, for NA=0.22 and *θ*_in_ = 0° the numerical results indicate that the centroid of the emission occurs at the taper’s tip with a standard deviation approximately zero, while the experimental results give a centroid distance from the tip of approximately 0.13 mm and a standard deviation of approximately 0.25 mm. However, both the numerical and the experimental intensity profiles for *θ*_in_ = 0° exhibit the maximum intensity at a distance of 0 mm from the tip (see Figs. 5a and 6a, respectively). Moreover, while the centroid distance obtained numerically increases linearly with the incident angle for the entire set of *θ*_in_, the experimental results deviate from a linear dependence at low input angles

At normal incidence, it can be analytically obtained that the mode guided with the largest fraction of the power (*l* = 0, *m* = 1) must reach the end of the fiber, as observed with the numerical calculations, because it becomes evanescent at a very small taper diameters (at 0.03 mm and 0.001 mm from the tip for the NA=0.22 and the NA = 0.66 fibers, respectively). This difference with respect to the experimental data, in which the emission region is more broadly spread along the taper (inset in Fig. 6a), can be explained considering that experiments are performed by launching light in a 1m-long patch cord, whose bends and length generate modal mixing. Energy from low order modes, and in particular from mode *l* = 0, *m* = 1, can be transferred toward modes with a marginally higher *k*_T_ [40, 41], generating a longer emission region in the experiments for low *θ*_in_.

Additionally, in the calculations we have assumed a perfectly linear taper profile and that both the core and the cladding of the fiber linearly decrease as one approaches the fiber tip in such a way that the ratio between the cladding and the core diameters is kept constant, but we cannot confirm that this is the case for the actual fiber. Similarly, the core and cladding of the actual fiber, which start being concentric at the input facet, could lose the symmetry as the fiber becomes thinner. Finally, all the assumed parameters (the refractive indexes and core and cladding diameters) were assumed as the nominal manufactures’ value, and the datasheet tolerances were not considered.

In conclusion, in this work we have presented a strategy to significantly reduce the computation cost and time required to calculate the wave propagation in long tapered optical fibers. We have applied this method to obtain the field distribution along tapered optical fibers of numerical aperture *NA* = 0.22 and *NA* = 0.66 assuming an incident Gaussian beam forming different angles with the fiber axis. Results have shown that the starting emission diameter and the intensity centroid distance from the tip of the fiber linearly increase with the light incident angle. We have also demonstrated that the calculation of a single mode is enough to estimate the starting emission diameter and the centroid distance from the tip, characterizing in this way the light delivery region as a function of the incident angle. Finally, we have validated the proposed strategy by comparing the obtained numerical and experimental results. Despite the paper is focused on pure dielectric taper, the method presented can be also applied as-is to metal-coated tapers. Compared to the ray tracing calculations usually employed for the theoretical characterization of tapered optical fibers, our technique gives access to the three components of the electric field along the fibers, which we believe can be exploited for designing novel generations of neural interfaces.

## Appendix A

### Comsol implementation

The beam-envelope method is implemented in Comsol Multiphyics by modifying the the variational weak form associated to the wave equation found in the *electromagnetic waves, frequency domain* module, which can be written as:

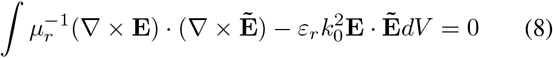

where 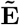 is the test function. If we substitute the electric field **E** in Eq. (8) by **E**′*e*^−*iβz*^, where **E**′ is the electric field of the slowly varying envelope function (in the *z* −direction) and *β* is the propagation constant of the input mode, the nabla operator can be transformed as.

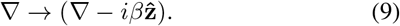

We modify the wave equation according to Eq. (9). That is, we included the extra terms containing −*iβ* to the expressions of the curl operator. More explicitly, for In cylindrical coordinates we have: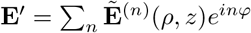,such that the curl operator in Eq. (9) becomes for each *n*:

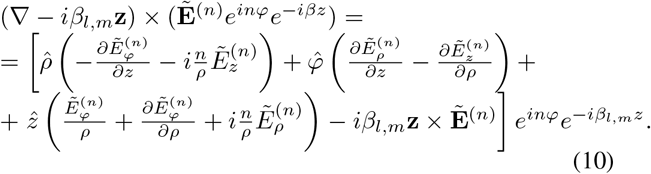

In this way, we solve the fields for **E**′ rather than the electric field **E**. After carrying out the calculations, we just need to multiply the results by *e*^−*iβz*^ in order to take into account the field oscillations due to the propagation constant *β*. Note that this method is implemented in the Optics module of Comsol Multiphysics only for trivial case when *n* = 0 (axisymmetric). Since we need to perform calculations with different values of *m*_az_, the manual implementation of the method was required. The axisymmetry space dimension can be selected in Comsol, which solves Eq. (4) for a specified value of *n*. The substitution in Eq. (9) must be implemented manually in the RF module. Perfectly matched layers (PMLs) are used in all the boundaries of the simulations, except for the input side for which we used ports.

## Acknowledgment

R.M-B., M.D.V., F. Pisanello and C.C. acknowledge funding from the European Union’s Horizon 2020 research and innovation program under grant agreement No 828972. F. Pisano and F. Pisanello acknowledges funding from the European Research Council under the European Union’s Horizon 2020 research and innovation program (grant agreement No 677683). M.D.V. acknowledges funding from the European Research Council under the European Union’s Horizon 2020 research and innovation program (No 692943). M.D.V. is funded by the US National Institutes of Health (U01NS094190). F. Pisanello and M.D.V. are funded by the US National Institutes of Health (1UF1NS108177-01).

